# Cereblon on Steroids: Beyond the Canonical Ligand Space

**DOI:** 10.64898/2026.08.28.747849

**Authors:** Alexander Herrmann, Christopher Heim, Samuel Maiwald, Iuliia Boichenko, Martin Neuenschwander, Andreas Oder, Birte Hernandez Alvarez, Andrei N. Lupas, Jens Peter von Kries, Marcus D. Hartmann

## Abstract

Cereblon (CRBN) is widely used in targeted protein degradation, but its ligand space has remained dominated by a narrow set of cyclic imide chemotypes. Here, we show that the accessible CRBN ligand space extends substantially beyond this canonical space. A high-throughput screen of > 40,000 compounds, followed by orthogonal biophysical validation, X-ray crystallography and SAR analyses, identified several chemically distinct ligand classes. These include linear acetyl-based motifs, a phthalide-derived scaffold, steroidal compounds, and a range of bicyclic ligands. They engage CRBN through distinct recognition modes, several of which deviate from the canonical hydrogen-bonding pattern. Steroidal scaffolds were particularly notable: cortisone binds the human CRBN thalidomide-binding domain with an affinity comparable to thalidomide, with its A-ring occupying the tri-tryptophan pocket in a glutarimide-like orientation despite lacking the canonical imide NH donor. SAR within this series showed substantial tolerance for chemical modification and scaffold simplification, raising the possibility that endogenous steroidal metabolites may contribute to the physiological ligand landscape of CRBN. Bicyclic lactams additionally provided synthetically accessible scaffolds with tunable affinity and promising sites for linker attachment. Across the identified ligand classes, none of the tested representatives induced detectable degradation of canonical CRBN neosubstrates, and several showed largely clean proteomic profiles. Together, these findings broaden the chemical, mechanistic and potential physiological landscape of CRBN recognition and provide diverse starting points for alternative, potentially neosubstrate-sparing CRBN recruiters.

## Introduction

Cereblon (CRBN) is a substrate receptor of a Cullin4-based E3 ubiquitin ligase complex and a central player in targeted protein degradation via the ubiquitin–proteasome system. Human CRBN (hCRBN) is a multi-domain protein with a C-terminal thalidomide-binding domain (TBD) that serves as the main substrate recognition site [1,2]. This domain harbors an aromatic cage formed by three conserved tryptophan residues (the tri-tryptophan pocket), which accommodates cyclic imide motifs through a conserved pair of hydrogen bonds involving the cyclic-imide carbonyl and NH groups [3,4]. The best-studied ligands of this pocket are thalidomide, related immunomodulatory drugs (IMiDs), and cereblon E3 ligase modulatory drugs (CELMoDs), which share a common binding mode centered on a five- or six-membered cyclic imide core [5], which is intrinsically susceptible to hydrolysis [6]. These compounds can act as molecular glues that promote the recruitment and ubiquitination of a set of neosubstrates, a process that depends on this canonical binding mode [7–12].

Only recently, C-terminal cyclic imide residues – specifically aspartimide and aminoglutarimide, which can arise spontaneously or be generated enzymatically – were identified as physiological ligands of CRBN [13–16]. Structurally, these natural degrons bind CRBN in the same canonical mode as IMiDs and related ligands, establishing CRBN as a receptor for cyclic imide motifs and linking endogenous and drug-induced substrate recognition [13]. Together, these findings show that CRBN recognizes a defined chemical motif, but leave open whether structurally more diverse moieties can engage the tri-tryptophan pocket with meaningful affinity and potentially reflect additional endogenous recognition modes.

In previous work aimed at delineating the chemical ligand space of the TBD, we established the single-domain homolog *Magnetospirillum gryphiswaldense* cereblon isoform 4 (MsCI4) as a versatile model system [4,5]. This protein proved robust and well suited for the development of a quantitative FRET assay amenable to high-throughput screening [17], as well as a crystal-soaking platform for high-resolution analysis of CRBN ligand binding modes [18]. Additionally, we developed a complementary microscale thermophoresis (MST) assay using the human TBD (hTBD) for quantitative affinity determination [19]. While these and related systems [20,21] enabled extensive exploration of thalidomide-derived scaffolds and related cyclic imides [22–27], the broader chemical space compatible with CRBN binding remains largely underexplored.

We therefore searched for CRBN ligands beyond canonical cyclic imides by combining an MsCI4-based FRET displacement assay with orthogonal validation by MST using the isolated hTBD and structural characterization in the MsCI4 crystal-soaking system. Applied to a chemically diverse library of > 40,000 small molecules, this workflow enabled broad exploration of CRBN-binding chemical space (Figure 1).

**Figure 1:**
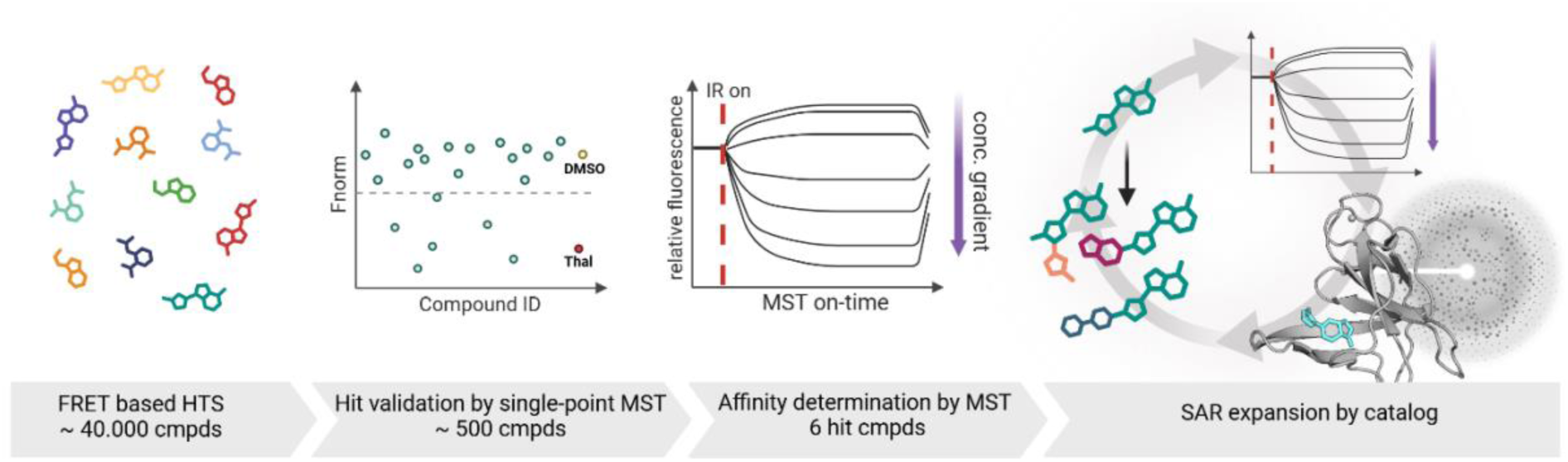
Description of the workflow.

Here, we report the outcome of this campaign and identify multiple structurally distinct CRBN-binding chemotypes beyond canonical cyclic imides, including scaffolds with unprecedented binding modes in the tri-tryptophan pocket. Among them, steroidal compounds emerged as an unexpected and particularly informative class, raising the possibility that endogenous steroidal metabolites may contribute to a broader physiological ligand landscape of CRBN. Collectively, these findings expand both the chemical and potential physiological landscape of CRBN recognition and provide new starting points for alternative CRBN recruiters.

## Results and Discussion

### High-throughput Screening

For the discovery of binding motifs beyond the canonical ligand space, we performed an HTS using the MsCI4-based FRET assay [17]. This assay employs *Magnetospirillum gryphiswaldense* cereblon isoform 4 (MsCI4) as a single-domain hTBD homolog, in which all strictly conserved residues involved in thalidomide binding are conserved, and a reporter ligand. This ligand, MANT-uracil, binds to the thalidomide-binding pocket with its uracil moiety, while its MANT portion forms a FRET pair with the tryptophans of the tri-tryptophan pocket. Accordingly, displacement of MANT-uracil by a test compound can be detected by loss of FRET signal, which we exploited in the HTS.

A total of 40,480 compounds from the FMP compound deck were screened using the FRET assay in a 384-well microtiter plate format. The compound deck encompassed commercially available compounds, including a diversity subset (28,512 compounds)[28], a fragment library subset containing smaller compounds with carboxyl and amine functional groups (4,576 compounds), and a subset of biologically annotated compounds (LOPAC, FDA-approved drugs, and drug candidates; 2,464 compounds). Another screened set consisted of non-commercially available compounds collected from academic research groups (4,928 compounds). The assay showed an average Z’-factor of 0.81 across the screened plates.

Of the 40,480 screened samples, 2,576 samples did not deliver a numeric readout on the plate reader due to signal overflow. Because of the short excitation wavelength of 295 nm, these samples exhibited autofluorescence and saturated the detector. A large number of compound samples led to increased fluorescence intensity at the MANT-uracil emission wavelength (Supplementary Fig 1A). Of the remaining samples, 646 showed a significant decrease in the MANT-uracil emission signal (Z-score less than -3). From these, the 352 samples with the lowest Z-scores were selected for an initial dose-response validation using the FRET assay.

When the picked compound plate was inspected, an enrichment of colored compounds was evident (Supplementary Fig 1B), indicating that colored compounds might interfere with the assay system—for example, by absorbing the excitation light and thereby decreasing the MANT-uracil emission intensity. To facilitate the detection of true FRET transfer (characterized by a decrease in MANT-uracil fluorescence only) during dose-response analysis, the emissions of both the MANT-uracil acceptor and the tryptophan donor were measured, and both values were normalized to the plate controls and plotted.

### Hit Validation

To overcome limitations arising from the MANT fluorophore, we employed an orthogonal microscale thermophoresis (MST) assay based on a BODIPY fluorophore for hit validation [19]. In this format, a BODIPY-uracil reporter was used together with the hTBD as an orthogonal protein construct. This assay has proven highly robust due to pronounced changes in the thermophoretic behavior of the BODIPY-uracil reporter depending on whether the reporter was in the hTBD-bound or free state. For select compounds, MST measurements were repeated using MsCI4 to assess species-specific differences.

From this point onward, we focus on four compounds and one additional ligand class that stood out because of their non-canonical chemistry (Figure 2): BS1, BS2, BS3, BS4, and the steroidal ligand class, represented here by cortisone. We describe the characterization of these ligands by crystallography, proteomics, and, where indicated, structure–activity relationship (SAR) analyses.

**Figure 2:**
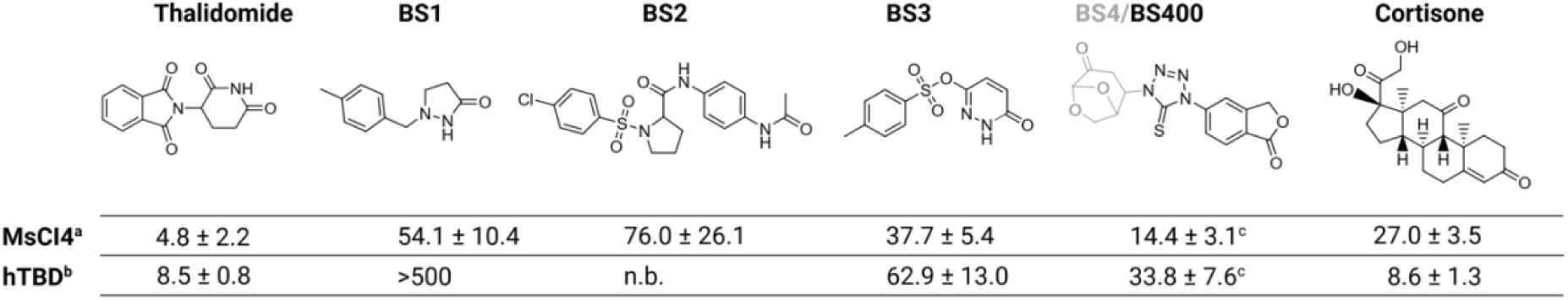
MST-based affinity determination of verified initial hits. ^a^ *K_i_* values determined by MST using NHS-RED labeled MsCI4. ^b^ *K*_i_ values calculated from IC_50_ values determined by competitive MST assay using hTBD and BODIPY-uracil as a reporter. Thalidomide is listed as a reference; all values are shown in µM (n=3). ^c^ Depicted K_i_ values are from BS400. The additional moiety present in BS4 (relative to BS400) is shown in grey. n.b. not binding.

### Characterization of BS1 and BS2

Compounds BS1 and BS2 stood out for their apparent species specificity, as they showed substantially higher affinity for MsCI4 than for hTBD (Figure 2). We performed a structural characterization in the MsCI4 soaking system for both compounds (Figure 3).

**Figure 3:**
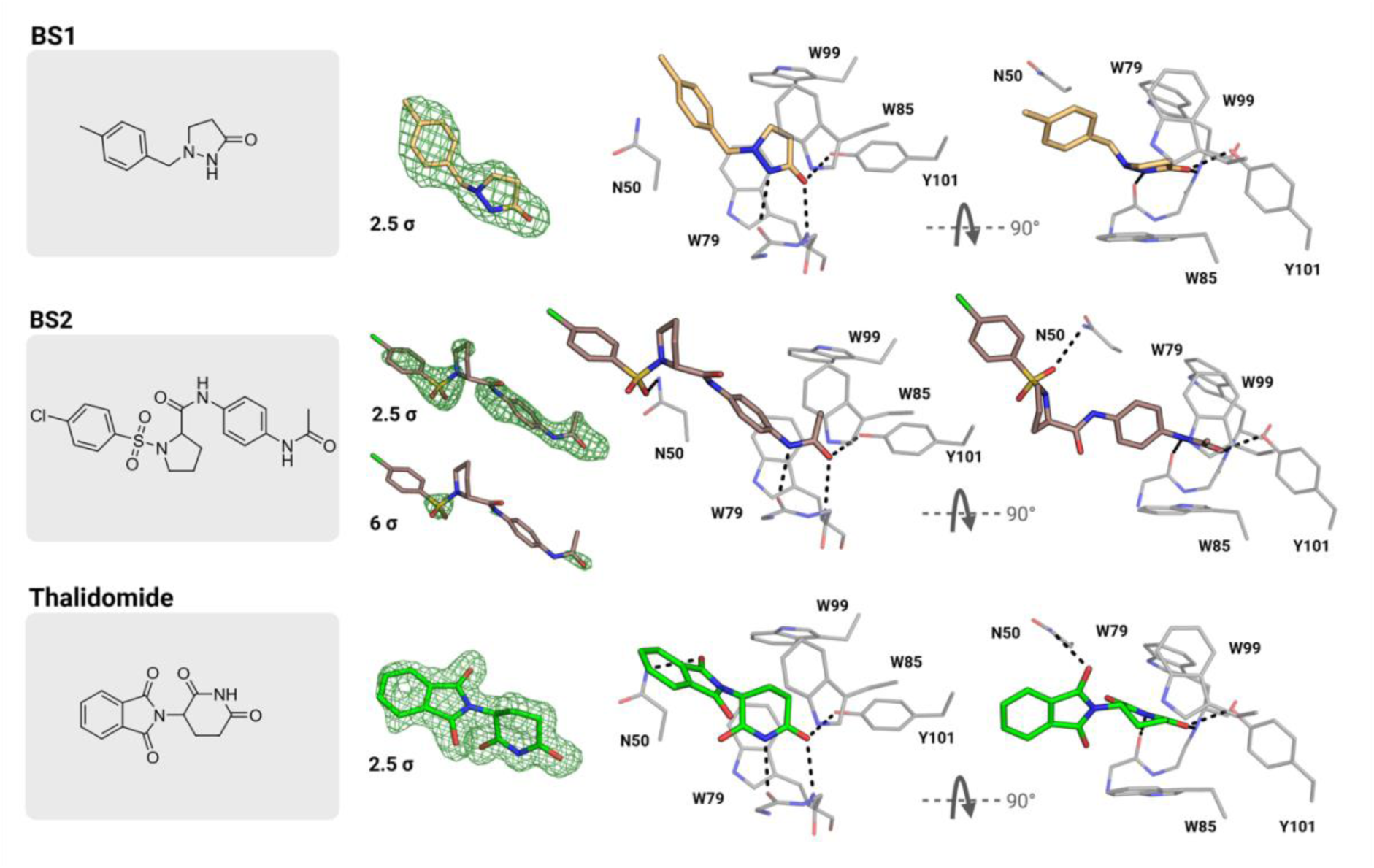
Binding modes of BS1 and BS2 in comparison to thalidomide. The compounds are shown with Fo-Fc omit maps contoured at the indicated contrast levels. The higher contrast level for BS2 identifies the sulfur. Hydrogen bonds are indicated with broken lines.

For BS1, the binding mode resembled that of γ-lactams [5]. Its pyrazolidin-3-one moiety forms the two strictly conserved hydrogen bonds within the tri-tryptophan pocket, in addition to a hydrogen bond to Y101 of MsCI4. In comparison to thalidomide, the most prominent difference lies in the shallower exit vector of the protruding moiety, which does not form additional H-bonding interactions.

For BS2, the acetyl group attached to the phenyl ring occupies the tri-tryptophan pocket in the same geometry as cyclic imides or lactams, forming the canonical pair of hydrogen bonds. To the best of our knowledge, this is the first instance of a linear binding motif observed for CRBN. Interestingly, the phenyl group is bound in an orientation perpendicular to that typically observed for cyclic imides, and the sulfonyl group forms a hydrogen bond to the asparagine in the sensor loop [29].

Since both compounds showed dramatically weaker affinity for hTBD than for MsCI4, we abstained from conducting further SAR analysis. The higher affinity for MsCI4 can be rationalized by the additional H-bond the compounds form with Y101 of the bacterial protein, which is one of the most prominent species-specific differences [2,4]. We propose, however, that such features could be exploited for the development of species-specific ligands. Furthermore, the linear binding mode of BS2 could hint at additional natural ligands, such as acetylated lysines, as previously hypothesized [5,30].

### Characterization of BS3

Compared to BS1 and BS2, BS3 stood out for its generally higher affinity for both MsCI4 and hTBD. Its crystallographic binding mode shares some similarities with those of BS1 and BS2 (Figure 4). Its pyridazin-3-one ring is bound in the tri-tryptophan pocket in a manner similar to BS1 and lactams in general, while its sulfonyl group forms an H-bond to the sensor-loop asparagine. The latter interaction is reminiscent of glutarimide-based sulfonyls that we had described previously [29].

**Figure 4:**
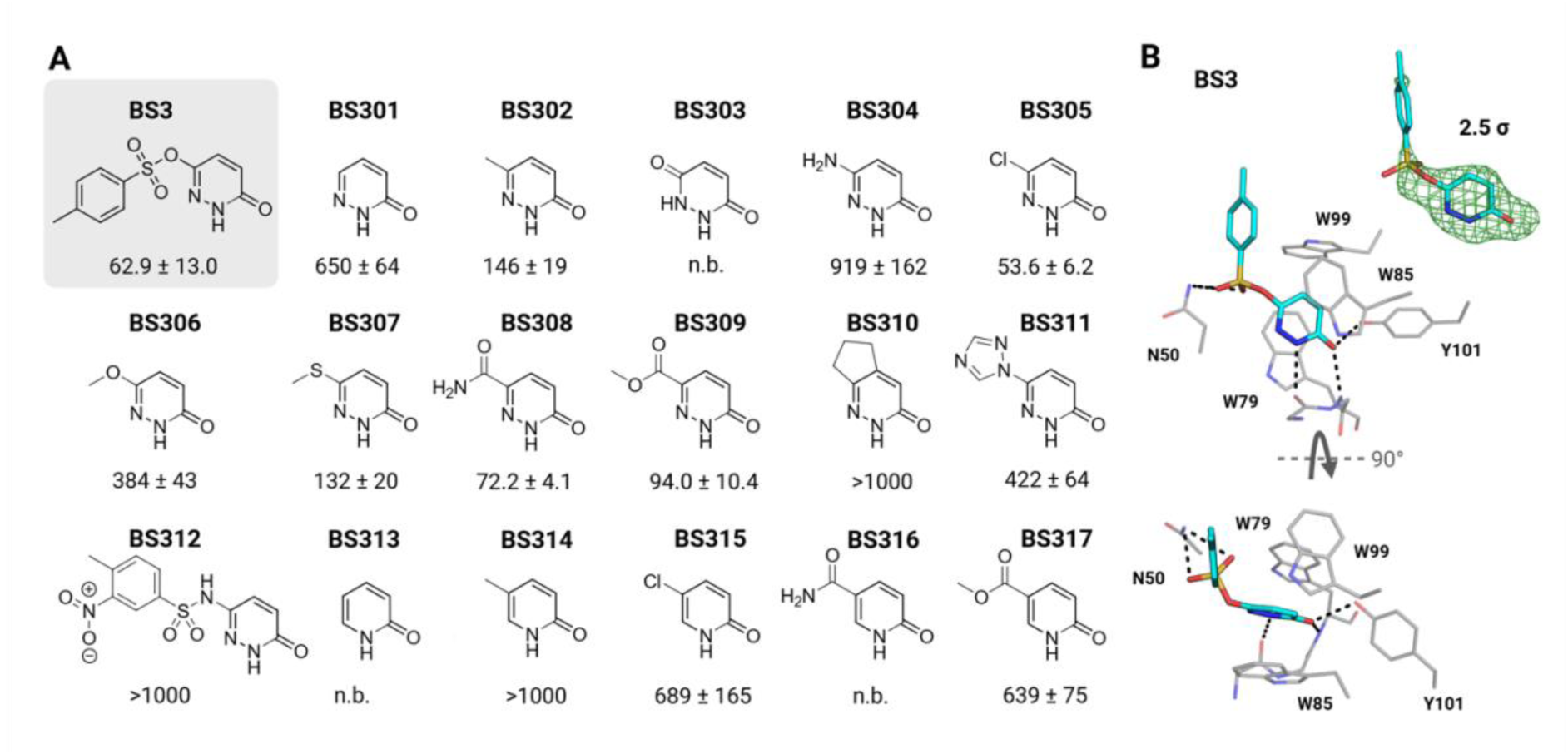
Binding mode and SAR of the BS3 scaffold. **A)** Affinities in µM for hTBD determined by MST. n.b. not binding; **B)** Binding mode of BS3 with Fo-Fc omit map contoured at 2.5 σ. Hydrogen bonds are indicated with broken lines.

Given its more favorable binding affinity, we performed an SAR analysis of 17 related compounds around this chemotype (Figure 4). Of these, however, only BS305 and BS308 matched the affinity of the initial BS3, indicating that the sulfonyl group and its interaction with the sensor-loop asparagine are key determinants of BS3 binding. A second consistent trend emerged from matched pyridazinone/pyridone pairs: replacement of the second ring nitrogen by carbon in BS313–BS317 consistently resulted in substantially weaker binding than for the corresponding pyridazinone analogues BS301, BS302, BS305, BS308, and BS309. This suggests that the second ring nitrogen contributes substantially to affinity, potentially by withdrawing electron density from the adjacent lactam NH and strengthening its canonical NH···O hydrogen bond within the tri-tryptophan pocket. Further optimization of the BS3 scaffold may therefore focus on derivatization of the distal sulfonyl substituent while retaining the pyridazinone core.

### Characterization of BS4

BS4 consists of a bridged bicycle linked via a tetrazole–5-thione moiety to a phthalide group. Therefore, it does not possess a cyclic imide or amide group as found in canonical binders. As we could not obtain the original compound for further characterization, we reverted to the simpler analog BS400, lacking the bridged bicyclic moiety. With a K_i_ value of 33.8 µM, BS400 has a higher affinity for hTBD than the tested BS1, BS2 and BS3 derivatives, indicating that the bridged bicycle is not necessary for binding.

To understand its binding mode, we employed the MsCI4 soaking system, revealing that the phthalide group occupies the canonical binding site, forming a hydrogen bond between the carbonyl and the W79 backbone NH, and another one between the ring oxygen and the Y101 side chain (Figure 5). Notably, the aromatic C7–H bond of the phthalide moiety points toward the W79 backbone carbonyl oxygen, with a C7···O distance of approximately 3.2 Å, a geometry compatible with a weak, non-classical C–H···O hydrogen bond [31]. This interaction may mimic the missing canonical imide-NH contact. In addition, the sulfur of the tetrazole-5-thione moiety forms an H-bond with the sensor-loop asparagine and participates in an additional water-mediated interaction at the pocket entrance. As Y101 is an MsCI4-specific feature, corresponding to F402 in hTBD, the phthalide group presumably forms only a single conventional H-bond within the tri-tryptophan pocket of the human protein, which may explain the higher affinity for MsCI4 (14.4 µM) as compared with hTBD (33.8 µM) (Figure 2). A limited SAR indicated that unsubstituted phthalide showed no appreciable affinity for hTBD. Among the 5-substituted analogues, the amino and cyano derivatives BS406 and BS408 showed notable affinity, with BS408 approaching that of BS400. Further polar substituents capable of engaging the sensor-loop asparagine, as observed for the tetrazole-5-thione moiety of BS400 and the sulfonyl group of BS3, may provide a route to improved affinity.

**Figure 5:**
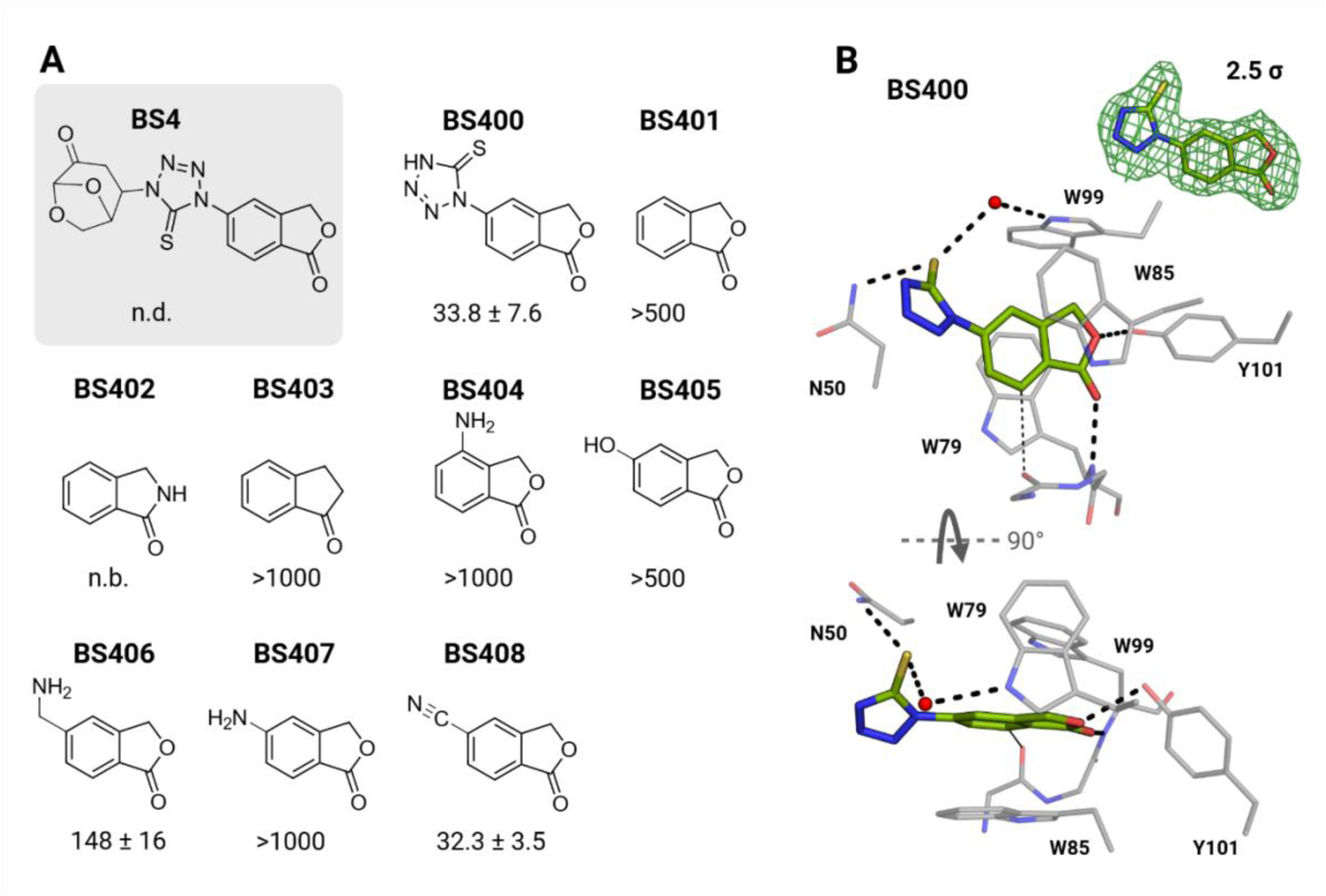
Binding mode and SAR of the BS4 scaffold. **A)** Affinities in µM for hTBD determined by MST. n.b. not binding; n.d., not determined. **B)** Binding mode of BS400 with Fo-Fc omit map contoured at 2.5 σ. The putative weak C–H···O hydrogen bond is indicated by a faint dotted line, other hydrogen bonds by broken lines.

### Characterization of steroidal ligands

We next investigated the binding mode of cortisone and related steroidal compounds, which lack the cyclic imide motif characteristic of canonical CRBN ligands. The MsCI4 soaking system revealed insertion of the steroid A-ring into the tri-tryptophan pocket in a geometry resembling that of glutarimide. However, the steroid scaffold lacks the imide NH donor that forms the canonical hydrogen bond to the W79 (W380 in human) backbone carbonyl. The vinylic C4–H bond of cortisone points toward this backbone carbonyl oxygen, with a C4···O distance of approximately 3.4 Å, a geometry compatible with a weak, non-classical C–H···O hydrogen bond analogous to the contact observed for BS400.

The A-ring carbonyl nevertheless forms the canonical hydrogen bond with the W79 backbone NH and an additional interaction with Y101 (Figure 6). Although the latter interaction is unavailable in hTBD, cortisone binds hTBD with even higher affinity than MsCI4, showing that the Y101 interaction is not required for high-affinity binding. In addition, the C-ring carbonyl forms a hydrogen bond with W99 at the pocket rim, mimicking an interaction observed in natural cyclic-imide degrons [13].

**Figure 6:**
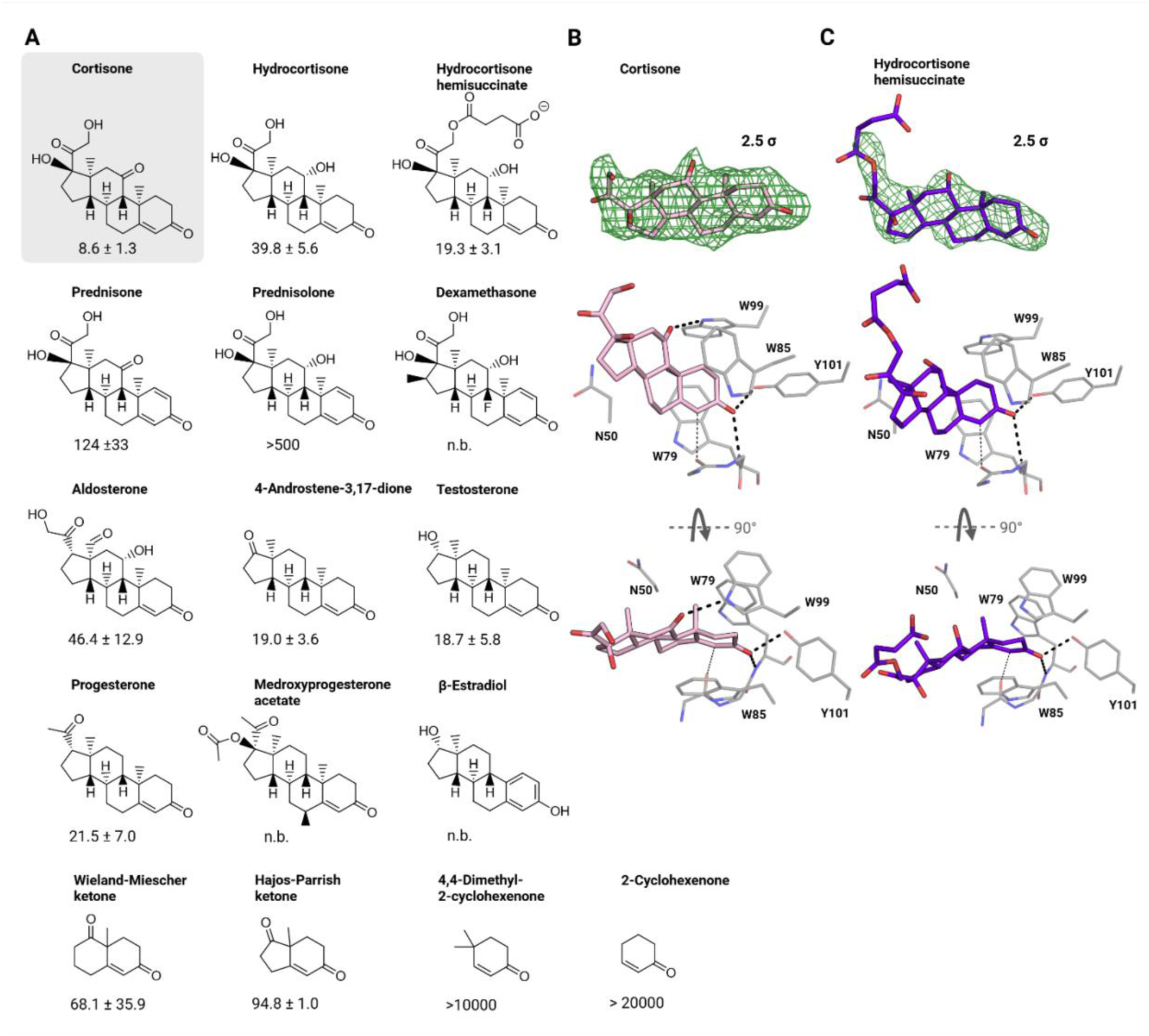
Binding mode and SAR of the steroid scaffold. **A)** Affinities in µM for hTBD determined by MST. n.b. not binding. **B)** and **C)** Binding modes of cortisone and hydrocortisone hemisuccinate with Fo-Fc omit maps contoured at 2.5 σ. The putative weak C–H···O hydrogen bond is indicated by a faint dotted line, other hydrogen bonds by broken lines.

A focused SAR across related steroidal and steroid-like scaffolds revealed several notable trends (Figure 6). First, introduction of an additional double bond in the A-ring, as in prednisone, reduced affinity. Second, replacement of the C-ring carbonyl by a hydroxyl group, as in hydrocortisone, abolished the corresponding hydrogen bond and consistently lowered affinity. Third, derivatives lacking the C-ring carbonyl or hydroxyl group, such as testosterone, nevertheless achieved affinities comparable to those of representatives with the C-ring hydroxyl group. Finally, substantial simplification of the steroidal scaffold, as exemplified by the Wieland–Miescher ketone, was tolerated.

### Characterization of bicyclic compounds

The steroidal series indicated that CRBN can accommodate rigid, polycyclic scaffolds and substantial variation outside the core-binding motif. We therefore explored bicyclic lactams as compact scaffolds combining conformational restriction with the canonical lactam hydrogen-bonding motif, while providing a second ring for further derivatization (Figure 7). Their binding modes resembled those of canonical ligands, with the lactam moieties forming the conserved pair of H-bonds within the tri-tryptophan pocket.

**Figure 7:**
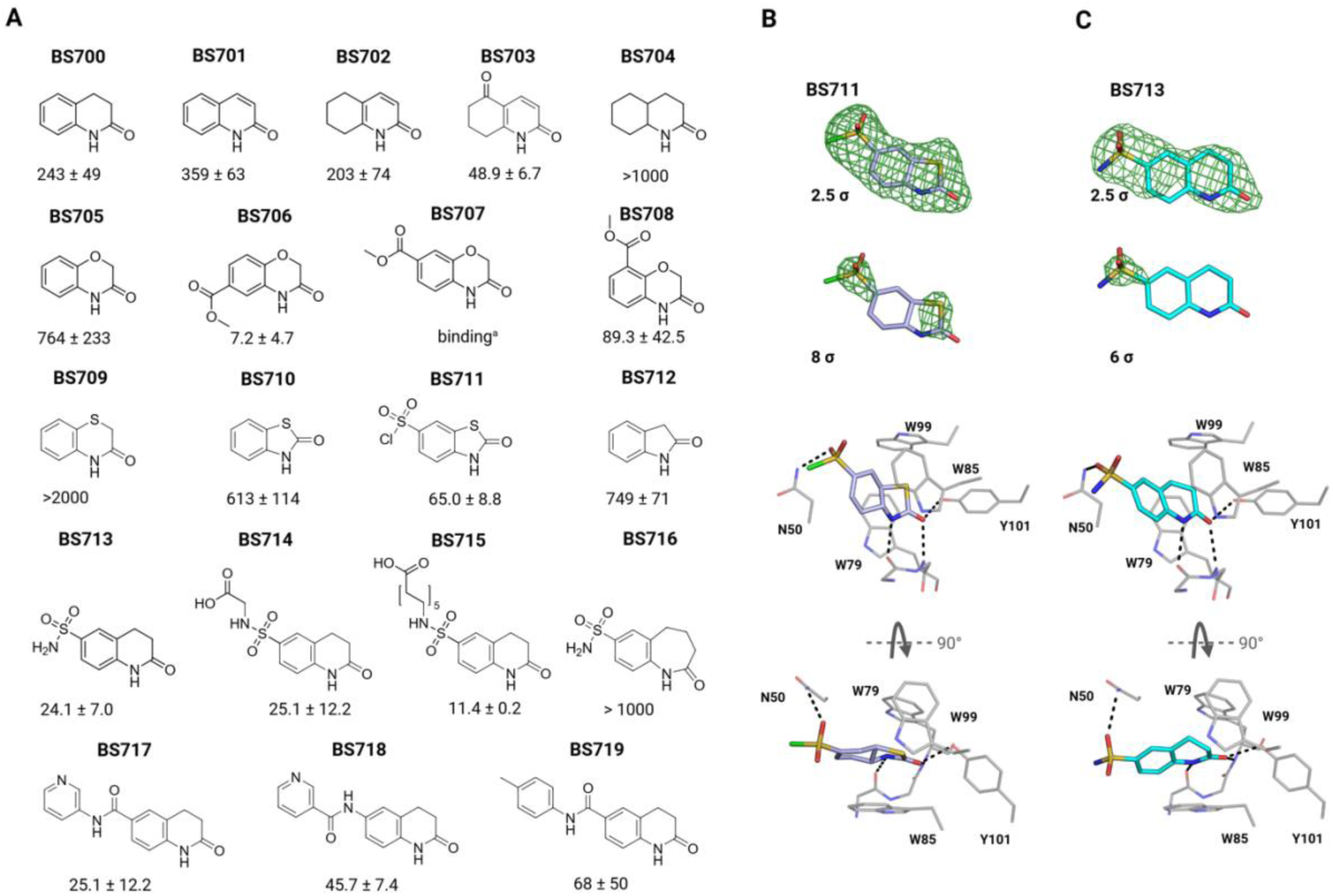
Binding mode and SAR of bicyclic compounds. **A)** Affinities in µM for hTBD determined by MST. ^a^ Binding was detected, but could not be quantified due to limited compound solubility. **B)** Binding mode of BS711 with Fo-Fc omit map contoured at 2.5 and 8 σ. **C)** Binding mode of BS713 with Fo-Fc omit map contoured at 2.5 and 6 σ. Hydrogen bonds are indicated with broken lines.

Variation of the core-binding ring showed that affinity was sensitive to both ring size and heteroatom composition. The six-membered lactam scaffold of BS700 retained binding upon incorporation of a double bond (BS701), whereas incorporation of oxygen or sulfur into this ring, as in BS705 and BS709, reduced affinity. Five-membered variants such as BS710 and BS712 showed only moderate affinity, while expansion to a seven-membered lactam, as in BS716 compared with BS713, was strongly detrimental.

The fused second ring proved more amenable to optimization. Aromatic and partially saturated variants retained binding, whereas complete saturation, as in BS704, was unfavorable. More substantial gains were achieved by substitution of the second ring: introduction of an additional carbonyl in BS703 improved affinity compared with BS702, while the methyl ester in BS706 increased affinity by more than two orders of magnitude compared with BS705. Several sulfonyl- and amide-containing derivatives likewise reached low- to mid-micromolar affinities. In the structures of BS711 and BS713, these gains can at least partly be rationalized by additional engagement of the sensor-loop asparagine (Figure 7). This SAR suggests that synthetically accessible bicyclic scaffolds can be optimized to exceed the affinity of cortisone. Furthermore, the tolerance of extended substituents in BS714 and BS715, with BS715 even showing increased affinity, identifies this region as a promising linker attachment site for bifunctional degraders.

### Neosubstrate engagement

With a set of new scaffolds in hand and their CRBN binding modes established, we next asked whether these ligands induce degradation of established CRBN neosubstrates. We therefore evaluated representative compounds in cell-based assays alongside established thalidomide derivatives as positive controls. The test set included BS1, BS2, BS3, and BS4, as well as representative steroidal and bicyclic compounds, and was assessed against IKZF3 [11,12], CK1α [32], and GSPT1 [33]. While the thalidomide-derived controls showed the expected depletion, none of the new compounds induced detectable degradation of the tested neosubstrates (Figure 8).

**Figure 8:**
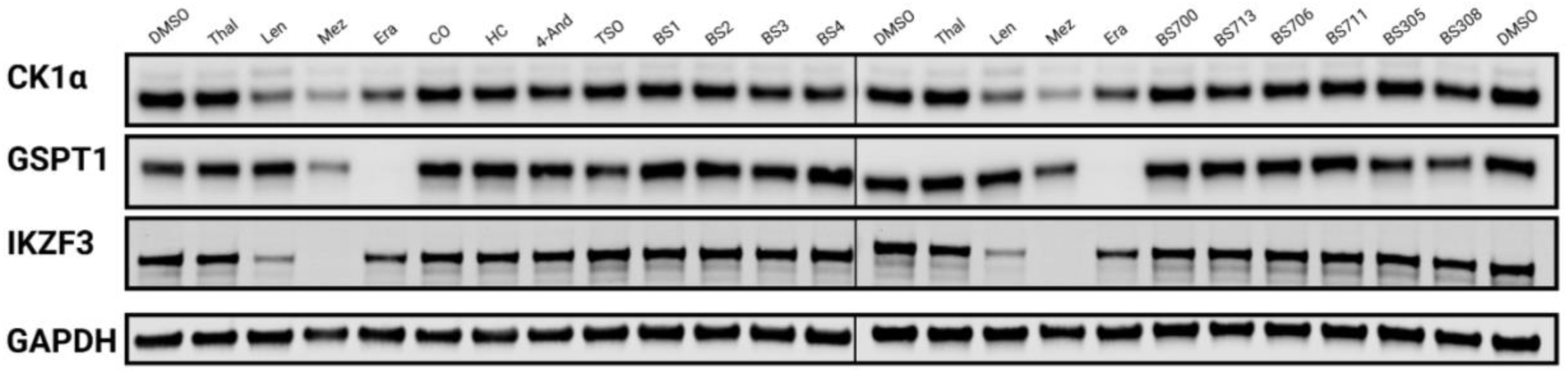
Western blots analysis of potential neosubstrate degradation. OPM-2 cells were treated with 10 µM of the indicated compounds for 24 h.The classical thalidomide (Thal) derivatives Lenalidomide (Len), Mezigdomide (Mez) and Eragidomide (Era) were used as a positive control. Compound abbreviations are defined as follows: CO, cortisone; HC, hydrocortisone; 4-And, 4-Androstene-3,17-dione; TSO, Testosterone.

To investigate whether the new ligands instead induce degradation of other, potentially novel CRBN neosubstrates, we next performed quantitative proteomic profiling for BS3, BS400, BS706, BS711, BS713, and cortisone (Figure 9). Consistent with the targeted assays, none of the compounds reduced the abundance of known CRBN neosubstrates. Moreover, BS3, BS400, and cortisone displayed largely clean proteomic profiles. In contrast, the bicyclic compounds BS706, BS711, and BS713 altered the abundance of several proteins, which may reflect indirect cellular effects or potential non-canonical substrate engagement requiring further investigation.

**Figure 9:**
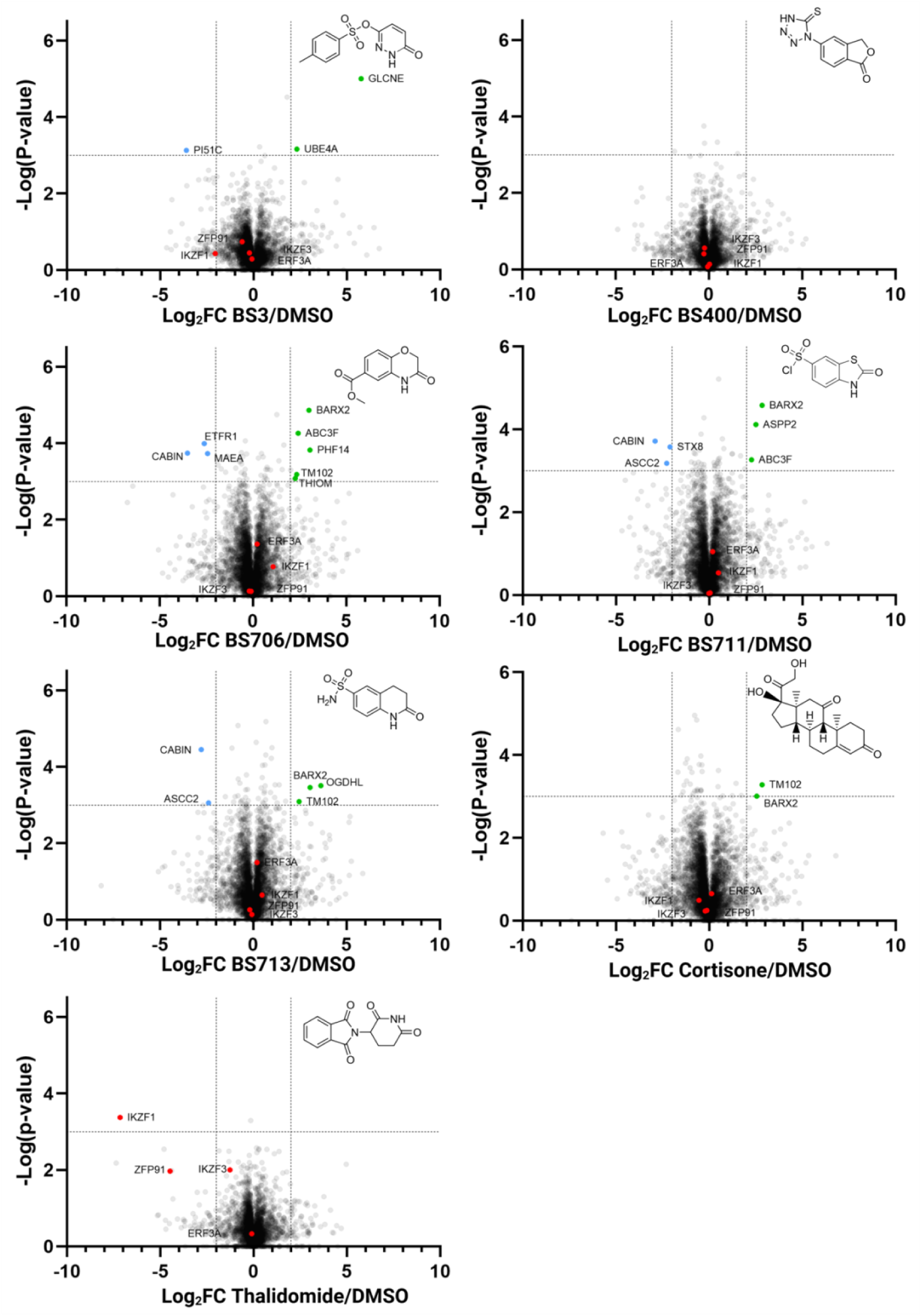
Quantitative proteomics for non-canonical CRBN binders. MM.1S cells were treated with compounds BS3, BS400, BS706, BS711, BS713, cortisone and thalidomide at 10 µM for 20 h. The identified proteins were plotted as log2 fold change (cmpd/DMSO) versus -log_10_ of the p-value. Proteins are considered to have significant change in abundance if -log_10_ (p-value) >3 and log2 fold change is >2 or < –2. This translates to a p-value < 0.001 and a fold change of 4 in abundance.

### Conclusions and outlook

CRBN ligands currently used in targeted protein degradation are dominated by a small set of cyclic imide motifs, most commonly glutarimide, uracil, or succinimide, with closely related exit vectors from the tri-tryptophan pocket and, in many cases, inherent hydrolytic liabilities. Our screening campaign reveals a substantially broader ligand space. We identified multiple chemically distinct ligand classes, including linear binding motifs and polycyclic scaffolds, that engage CRBN through alternative recognition modes, several of which do not reproduce the full canonical hydrogen-bonding pattern of established binders. Among these, BS2 is of particular conceptual interest, as it provides the first verified example of a linear acetyl-based motif occupying the core-binding pocket of CRBN. In this case, productive binding is supported by auxiliary interactions outside the canonical pocket. Although its pronounced preference for MsCI4 limits direct transferability to the human protein, the finding nevertheless raises the possibility that acetyl groups, such as in post-translational modifications, could be recognized by CRBN in an appropriate sequence or structural context, thereby pointing to potential physiological recognition principles beyond C-terminal cyclic imides [5].

BS3 and BS4 extend the CRBN ligand space in different directions. BS3 retains the canonical hydrogen-bonding pattern, while the matched pyridazinone/pyridone SAR suggests that lactam-based CRBN binding can be tuned through heterocycle design, with pyridazinone emerging as a particularly favorable motif. BS4, by contrast, retains only one of the two canonical H-bonds within the tri-tryptophan pocket, while a putative weak C–H···O interaction may mimic the missing imide-NH contact. The scaffold nevertheless retained appreciable affinity for hTBD, identifying the phthalide-based core as an alternative CRBN-binding motif with potential for distinct exit vectors.

Steroidal scaffolds emerged as the most striking non-canonical ligand class. Without further optimization, cortisone displayed one of the highest affinities for hTBD in this study, comparable to that of thalidomide, despite lacking the cyclic imide motif. Its A-ring occupies the tri-tryptophan pocket in a glutarimide-like orientation, while the scaffold lacks the canonical imide NH donor. Instead, the corresponding C4–H bond adopts a geometry compatible with a weak non-classical C–H···O interaction with the same backbone carbonyl, analogous to the contact observed for BS400. In addition, steroids containing a C-ring carbonyl can establish an additional interaction at the rim of the tri-tryptophan pocket that resembles contacts observed for natural degron peptides [14]. The expanded SAR shows that CRBN binding is retained across structurally diverse steroidal compounds and substantially simplified steroid-like scaffolds, while not being a general property of steroids. These observations raise the possibility that endogenous steroidal metabolites may represent an overlooked component of the physiological CRBN ligand landscape.

Finally, the steroidal compounds inspired the exploration of bicyclic lactams as compact, synthetically accessible CRBN ligands. SAR indicated a preference for a six-membered core-binding lactam, whereas the fused second ring tolerated extensive modification and offered substantial scope for affinity optimization, with the best compounds competing with the affinity observed for cortisone. Extended substituents were also well tolerated, providing a promising linker attachment site.

None of the tested representatives induced detectable degradation of the canonical CRBN neosubstrates examined here, and several compounds showed largely clean proteomic profiles. Together, these findings broaden both the chemical and potential physiological landscape of CRBN recognition and provide structurally diverse starting points for alternative, potentially neosubstrate-sparing CRBN recruiters. By providing alternative exit vectors and scaffold geometries, these ligands may enable CRBN-based degraders that form ternary complexes inaccessible to classical IMiD- and CELMoD-based PROTACs and molecular glues, thereby broadening the range of efficiently degradable targets.

## Methods

### Protein expression and purification

***Wildtype MsCI4*** was cloned, expressed and purified as described previously [17]. The MsCI4 construct in the pETHis1a vector was transformed into *E. coli* C41 (DE3) cells. Cultures were grown in LB medium supplemented with 50 µM ZnCl₂ at 37 °C until reaching an optical density (OD₆₀₀) of 0.6, protein expression was induced by the addition of 1 mM IPTG and performed for 4 h at 37°C.

The purification-buffer used during purification contains 20 mM TRIS pH 8, 0.15 M NaCl and 5 mM β-Mercaptoethanol. During lysis EDTA-free c0mplete protease inhibitor and DNase I was supplemented. After loading the Ni-NTA matrix, the column was washed with the buffer containing 50 mM imidazole and 100 mM subsequently. The tagged protein elutes at 200 mM imidazole.

The eluted protein was dialyzed in purification-buffer (change buffer once) in the presence of TEV protease. Reverse Ni-NTA affinity purification was then performed to remove the cleaved tag and TEV protease, with MsCI4 eluting at 70 mM imidazole.

***hTBD (residues 319-425) with a C366S*** point mutation was cloned into pET24a with a His-Sumo Tag. Expression was done in *E. coli* BL21 (DE3) [19]. The cultures were grown in TB medium supplemented with kanamycin and 50 µM ZnCl2. Upon reaching an optical density (OD_600_) of 0.6, protein expression was induced by adding 1 mM IPTG. Expression takes place over night at 16°C.

The buffer used for affinity chromatography contains 50 mM TRIS pH 8.0, 500 mM NaCl, 10 µM ZnCl2, 1 mM TCEP and 20 mM imidazole. This condition is also used for lysis with the addition of 0.1 % (v/v) Triton X-100 and c0mplete. After loading the His-trap, the column was washed properly with 5 column volumes of buffer. By adding 480 mM imidazole to the buffer, elution was done in a gradient using both buffer compositions. The His-Sumo fused hTBD elutes at approximately 120 mM imidazole. The pooled fractions were dialyzed for 2-steps over night with ULP1-protease (1:100), using the standard buffer with 20 mM imidazole. Reverse Ni-NTA affinity purification was performed to remove the tag and the tagged protease.

### FRET-based HTS

MANT-uracil was obtained by custom synthesis (Jena Bioscience) and prepared as a 1 mM stock solution in DMSO. All high-throughput screening experiments were conducted in FRET buffer (20 mM Tris/HCl, pH 7.5, 150 mM NaCl, 0.5 mM β-mercaptoethanol, 0.1% Tween20) in a final reaction volume of 20 µl in the presence of 3 µM MsCI4 and 3 µM MANT-Uracil.

#### Primary screening

First, 10 µl of FRET buffer were dispensed into a white low-volume 384-well microplate (ProxiPlate 384-shallow well, Revvity, #6008280) using a Biotek MultiFlo dispenser (Agilent) equipped with a 5 µl dispensing cassette. Compounds were transferred from compound mother plates (containing compounds dissolved to 10 mM in DMSO in columns 1–22, and DMSO alone in columns 23 & 24) to the assay plates using an FP1 pin-tool (V&P Scientific) mounted in a liquid handling workstation (FxP, Beckman Coulter). Using dyes dissolved in DMSO we determined the average transfer volume of the Pin-Tool being around 20 nL. MsCI4 and MANT-uracil solutions were prepared at 2x final concentration and incubated for 30 minutes at room temperature. Using the microplate dispenser, 10 µl of the 2x MsCI4/MANT-uracil solution was dispensed into columns 1–23 of the assay plate. Using a manual multi-pipette, 10 µl of 2x MANT-uracil solution was added to column 24. Thus, compound samples were present at an approximate concentration of 10 µM in columns 1–22; the MsCI4/MANT-uracil solution receiving only DMSO in column 23 served as the 100% binding control, and the MANT-uracil solution receiving only DMSO in column 24 served as the 0% binding control.

The assay plates were incubated for one hour at room temperature and then read on a microplate reader (Safire2, Tecan) with the following settings: monochromators set to an excitation wavelength of 295 nm and an emission wavelength of 440 nm (bandwidth 9 nm), reading top fluorescence with 10 flashes. The data were normalized for each plate; %Activity values for each sample were determined by normalizing to the medians of the 100% and 0% reference samples on the plate. Z-score values (distance to the mean in units of standard deviation) were calculated using all compound samples of a plate in columns 1–22. Z’-factors were calculated using the 100% and 0% control samples of each plate [34].

#### Dose-response measurements

For dose-response measurements, compounds were picked as 10 mM DMSO solutions onto a new compound mother plate. This compound mother plate was then serially diluted across plates by adding 5 µl of DMSO, transferring 5 µl of the mixture to a new empty microplate, and repeating this step multiple times. This created a dilution series with 9 consecutive 2-fold dilutions, with 5 mM as the highest compound stock concentration.

Using these prediluted compound mother plates, each dilution was assayed in two technical replicates using the same protocol as the primary screening, with the following changes: the pin-tool transfer was performed three times for each plate, resulting in a concentration range of 120 nM to 30 µM. Alongside the 440 nm emission (MANT-uracil acceptor fluorescence), the emission at 360 nm (Tryptophan donor fluorescence) was measured as well. Dose–response curves and half-maximal inhibitory concentration values were determined by non-linear regression using a four-parameter logistic (4PL) model in Pipeline Pilot (version 8.5).

### Microscale thermophoresis (MST)

#### Labeled MST

MsCI4 was covalently labeled using the Monolith Protein Labeling Kit RED-NHS 2nd Generation (NanoTemper Technologies, MO-L011) according to the manufacturer’s protocol with the following modifications. To increase labeling yield, 30 µM protein was used for the buffer exchange step. As MsCI4 binds DMSO, the labeling dye was reconstituted in labeling buffer. Following the labeling reaction, unbound dye was removed by gravity-flow SEC equilibrating the column with MST buffer (50 mM Tris pH 7.4, 150 mM NaCl, 10 mM MgCl₂, 0.05% Tween-20, 1 mM TCEP). Fractions with protein concentrations greater than 2 µM and a degree of labeling of 0.5–1.0 were pooled, aliquoted, and flash-frozen in liquid nitrogen. Labeled MsCI4 was used at a final concentration of 20 nM in MST measurements.

#### Competitive MST

Binding affinities (K_d_) of Uracil-based reporter ligands to hTBD were determined by microscale thermophoresis (MST) using a 16-point 1:1 dilution series against 200 nM reporter (as described previously [19]). The reporter was prepared from a 2 mM DMSO stock, diluted 1:50 in MST buffer (50 mM Tris pH 7.4, 150 mM NaCl, 10 mM MgCl₂, 0.05% Tween-20, 1 mM TCEP) to 40 µM, protected from light, and kept on ice. For measurements, this stock was further diluted 1:100 and mixed 1:1 with hTBD dilution series to achieve 200 nM final reporter concentration.

For competition assays, compounds were prepared as 16-point 1:1 DMSO dilution series, each point further diluted 1:100 in water, and mixed with a protein–reporter complex prepared with a protein concentration at twice the previously determined K_d_ and 400 nM reporter. Mixing 1:1 with diluted compounds resulted in a final DMSO concentration of 0.5% (v/v).

Measurements were performed on a Monolith NT.115 (NanoTemper Technologies) using standard capillaries (red channel, 20% LED power, medium MST power) in at least duplicates for binders.

Data were analyzed in GraphPad Prism v9.5.0 using a four-parameter dose–response fit. IC₅₀ values were converted to K_i_ using the Cheng–Prusoff equation [35].

### Cell culture

MM.1S and OPM-2 cells were cultured at 37°C in a humidified incubator with 5% CO2. Cells were maintained in RPMI1640 medium supplemented with 10% fetal calf serum and 1% penicillin-streptomycin. Cells were split every 2-3 days to avoid overgrowth. As MM.1S formed confluent subculture, a cell scraper was used to detach cells prior to splitting.

### Western blotting

OPM-2 cells were seeded at ∼0.6 × 10⁶ cells/mL one day prior to treatment. Cells were exposed to compounds (10 µM, 0.1% DMSO) for 24 h, harvested by centrifugation, washed with ice-cold PBS, flash-frozen in liquid nitrogen, and stored at −80 °C.

For lysis, pellets were resuspended in lysis buffer (20 mM Tris pH 7.7, 150 mM NaCl, 1 mM EDTA, 1 mM EGTA, 1% Triton X-100, 2 mM MgCl₂, supplemented with protease inhibitors and DNase I) and incubated on ice for 30 min. Lysates were cleared by centrifugation and protein concentration was determined by BCA assay.

Samples were prepared in LDS sample buffer with 5% β-mercaptoethanol, heated at 70 °C for 10 min, and separated on 4–12% Bis-Tris gels. Proteins were transferred to nitrocellulose or PVDF membranes using a semi-dry transfer system. Membranes were blocked with 5% milk in TBS-T and incubated with primary antibodies overnight, followed by HRP- or fluorophore-conjugated secondary antibodies.

Primary antibodies: Aiolos (Rabbit mAb, CST #15103, D1C1E, 1:2000), CK1α (Rabbit mAb, Abcam #ab206652, 1:2000), eRF3/GSPT1 (Rabbit mAb, CST #14980, 1:2000), GAPDH (Rabbit mAb, CST #2118, 14C10, 1:2000)

Secondary antibodies: Peroxidase AffiniPure® Goat Anti-Rabbit IgG (H+L) (Jackson ImmunoResearch), used with 5% milk in TBS-T; IRDye 800CW Goat Anti-Rabbit (LI-COR), used with 5% BSA in TBS-T.

### Proteomics

MM.1S cells were seeded one day in advance at ∼0.6 × 10⁶ cells/mL in 6-well plates (4 mL per well). Cells were treated with compounds (10 µM, 0.1% DMSO) for 20 h, harvested by centrifugation (1000 g, 5 min), and washed twice with ice-cold PBS. For proteomics, cell pellets were snap-frozen in liquid nitrogen and shipped on dry ice to Biogenity ApS (Aalborg, Denmark), where lysis, digestion, LC–MS/MS analysis, and primary processing were performed using standard workflows. Raw data were returned and analyzed in Perseus v2.0.11 [36]. Data were log_2_-transformed, normalized (width adjustment), and filtered for proteins with ≥85% valid values. Statistical analysis and visualization were performed in GraphPad Prism v9.5.0, including volcano plots of log_2_ fold change versus −log(p-value) relative to DMSO control.

### MsCI4 crystallization

The crystals used in the soaking experiments were obtained as described previously [18]. MsCI4 was concentrated to 17 mg/mL. As precipitant, (NH₄)H₂PO₄ at concentrations between 0.4 and 0.6 M was used. In the presence of 3 mM thalidomide and mixed 1:1 (v/v) with the precipitant solution, crystals grew within a few days. For soaking, crystals were transferred into drops of fresh reservoir solution supplemented with the desired compound, either from DMSO stock or as dried compound. After 24 h, this step was repeated by transferring the crystals into fresh drops to ensure complete exchange of thalidomide. For cryoprotection, sodium malonate was added to a final concentration of 70% to the reservoir solution prior to flash-cooling in liquid nitrogen.

### Crystallographic Data collection, processing and structure determination

Diffraction data were collected at beamlines X10SA (SLS) and ID23-1 (ESRF). Data were processed using XDS [37] . Structures were solved by molecular replacement using PDB entry 4V2Y as the starting model and refined with REFMAC5 [38]. Iterative model building and real-space refinement were performed in Coot v0.9 [39]. Molecular graphics were generated using PyMOL Molecular Graphics System, version 3.1 [40]. Data collection and refinement statistics are summarized in Tables S1 and S2.

## Acknowledgements

We thank Reinhard Albrecht and Luca Bischof for assistance with crystallographic data collection and processing, and the staff of beamline X10SA at the Swiss Light Source (Paul Scherrer Institute, Villigen, Switzerland) as well as the staff of beamline ID23-1 at the European Synchrotron Radiation Facility (ESRF, Grenoble, France) for excellent technical support. This work was supported by institutional funds from the Max Planck Society.

