## Supplemental Information for "Cereblon on Steroids: Beyond the Canonical Ligand Space"

**Contents:**

| | |
|---|---|
| MsCI4-based FRET assay readout and scatter chart | page 3 |
| Binding mode of BS305 & BS308 | page 4 |
| Binding mode of BS700 | page 4 |
| Neo-substrate degradation assay western blots | page 5 |
| Crystallographic data collection and refinement statistics | pages 6-7 |

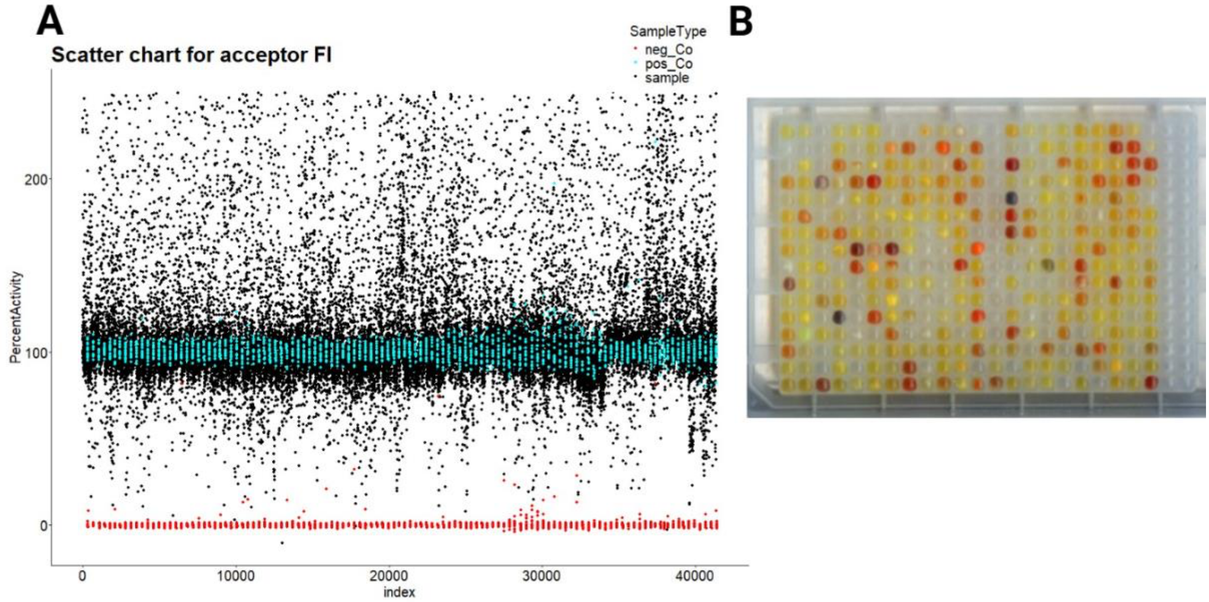


**Figure S1: MsCl4-based FRET assay readout (A)** Scatter diagram of %activity values of all measured compound samples, displaying the normalized readout at 440 nm (MANT-uracil emission). 100% refers to the fluorescence intensities observed in reference samples containing MsCl4/MANT-uracil (cyan), while 0% refers to MANT-uracil alone (red). Evident are numerous samples delivering fluorescence intensities above those of bound MANT-uracil (points above the cyan dots), pointing to compound autofluorescence. Primary hits show decreased MANT-uracil intensities between the 100% and 0% controls. **(B)** Scan of the compound mother plate containing the 352 most active samples after primary screening. Evident is the strong enrichment of colored samples.

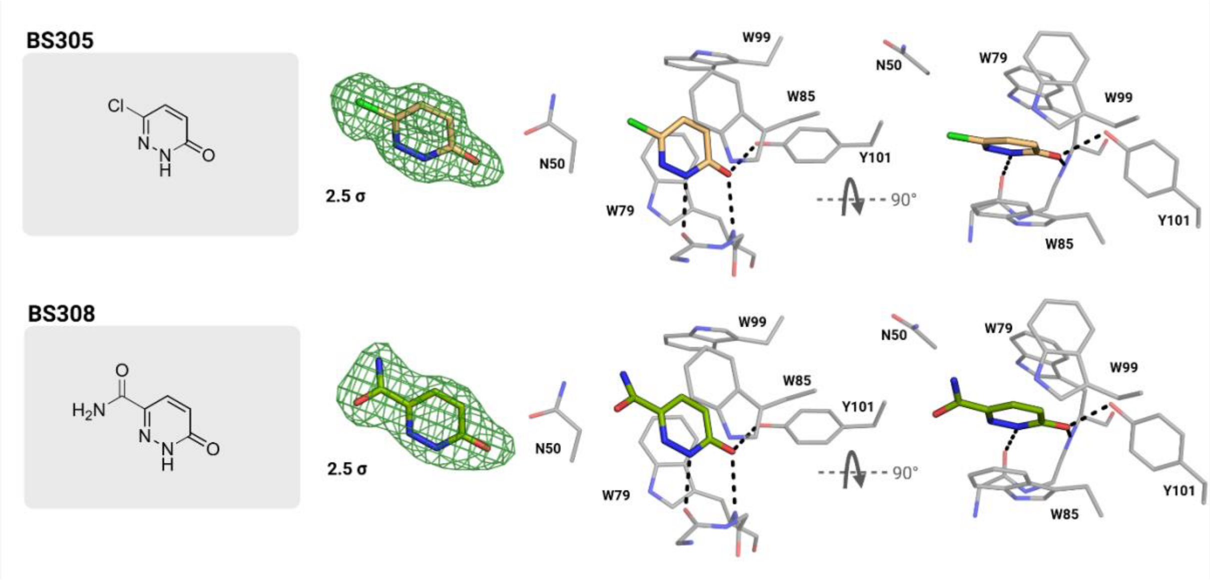


**Figure S2: Binding mode of BS305 and BS308** with Fo-Fc omit map contoured at 2.5 σ. Hydrogen bonds are indicated with broken lines.

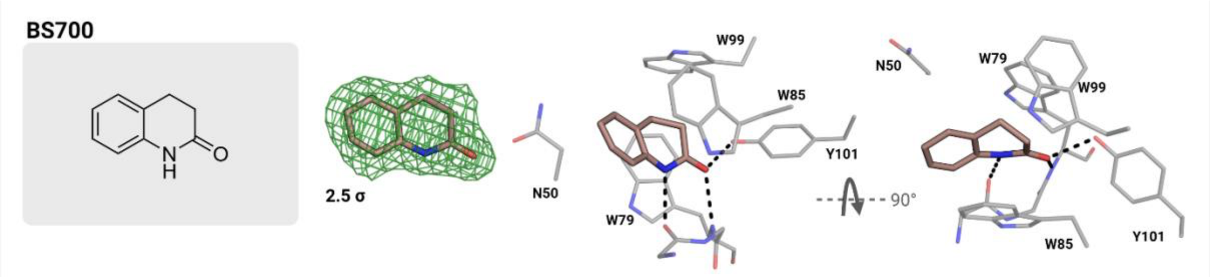


**Figure S3: Binding mode of BS700** with Fo-Fc omit map contoured at 2.5 σ. Hydrogen bonds are indicated with broken lines.

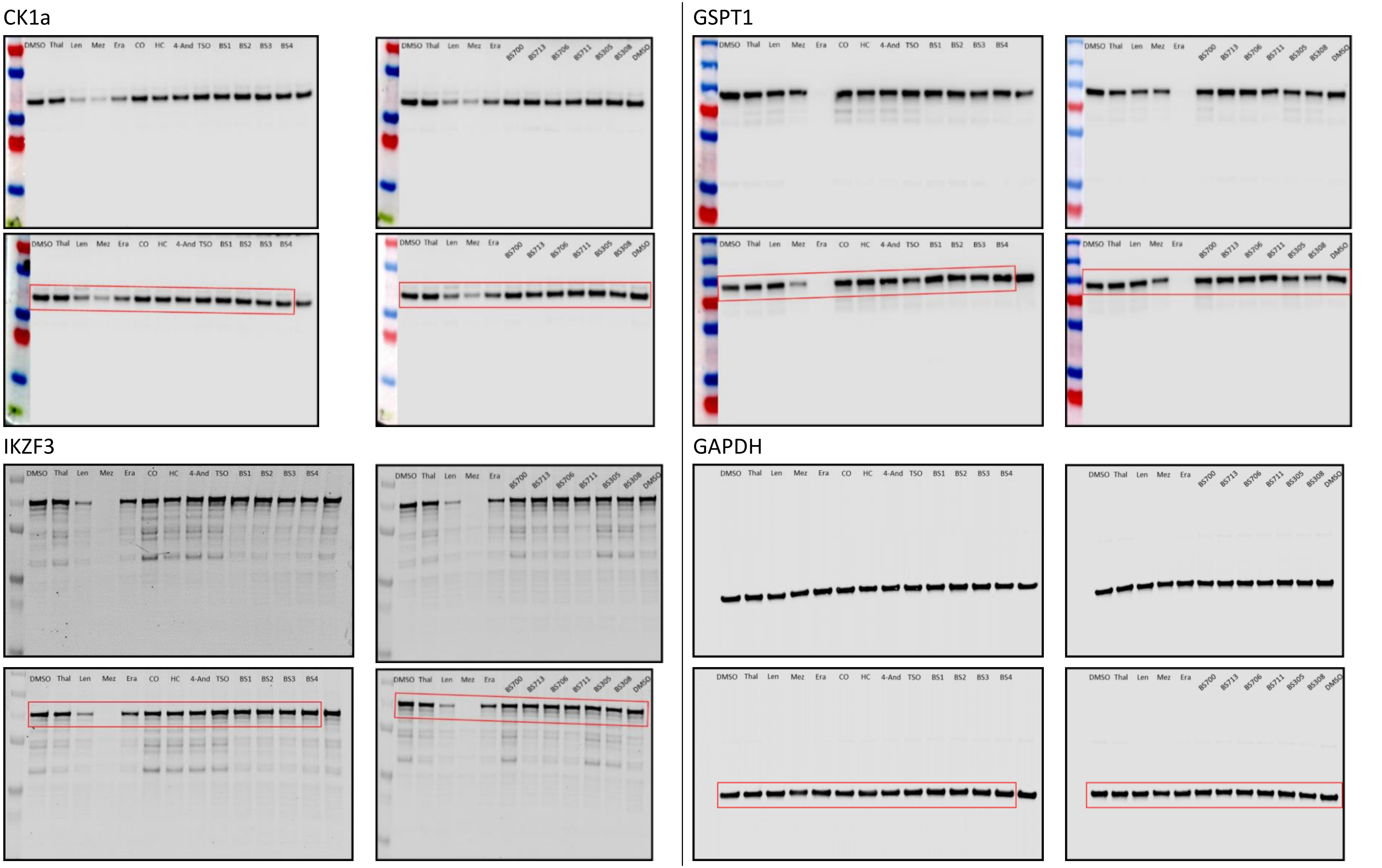


**Figure S4: Neo-substrate degradation assay.** OPM-2 cells were treated with 10 µM of the indicated compounds for 24 h to assess potential neo-substrate degradation of CK1a, GSPT1 and IKZF3. Analyzed by Western blot in biological duplicates. The classical thalidomide (Thal) derivatives Lenalidomide (Len), Mezigdomide (Mez) and Eragidomide (Era) were used as a positive control. Compound abbreviations are defined as follows: CO, cortisone; HC, hydrocortisone; 4-And, 4-Andristene-3,17-dione; TSO, Testosterone. The red-marked regions indicate the blots shown in the main manuscript (Figure 8).

**Table S1: Data collection and refinement statistics** for the crystallographic datasets of MsCI4 with bound BS1, BS2, BS3 and its derivatives BS305 and BS308, and BS400. Values in parentheses refer to the highest resolution shell

| | **BS1** | **BS2** | **BS3** | **BS305** | **BS308** | **BS400** |
|---|---|---|---|---|---|---|
| **PDB accession code** | **34BS** | **34BT** | **34BU** | **34BV** | **34BW** | **34BX** |
| Wavelength (Å) | 1.000 | 0.886 | 1.000 | 1.000 | 1.000 | 1.000 |
| Resolution range (Å) | 37.38 -2.09 (2.17 - 2.09) | 44.12 -1.7 (1.76 -1.7) | 34.92 -1.83 (1.90 -1.83) | 36.98 -1.44 (1.49 -1.44) | 49.28 -1.9 (1.97 -1.9) | 41.46 -1.91 (1.98 -1.91) |
| Space group | $P2_12_12_1$ | $P2_12_12_1$ | $P2_12_12_1$ | $P2_12_12_1$ | $P2_12_12_1$ | $P2_12_12_1$ |
| a, b, c (Å) | 56.95, 59.97, 87.99 | 56.43, 60.13, 88.23 | 57.24, 59.53, 88.13 | 56.79, 58.99, 86.44 | 56.59, 59.29, 88.59 | 57.04, 60.36, 88.43 |
| α, β, γ (°) | 90.0, 90.0, 90.0 | 90.0, 90.0, 90.0 | 90.0, 90.0, 90.0 | 90.0, 90.0, 90.0 | 90.0, 90.0, 90.0 | 90.0, 90.0, 90.0 |
| Total reflections | 240951 (22345) | 443228 (46011) | 323012 (16387) | 691653 (63443) | 317640 (32681) | 306322 (20579) |
| Unique reflections | 18442 (1804) | 33730 (3322) | 26933 (2445) | 53099 (5231) | 24148 (2383) | 24277 (2396) |
| Multiplicity | 13.1 (12.4) | 13.1 (13.9) | 12.0 (6.7) | 13.0 (12.1) | 13.2 (13.7) | 12.6 (8.6) |
| Completeness (%) | 99.96 (100.00) | 99.98 (100.00) | 98.88 (91.54) | 99.90 (99.54) | 99.96 (100.00) | 99.86 (99.62) |
| Mean I/σ (I) | 13.25 (1.47) | 14.5 (1.22) | 18.68 (1.16) | 19.97 (1.15) | 8.34 (1.11) | 17.58 (1.11) |
| Wilson B-factor | 41.12 | 30.29 | 35.50 | 23.44 | 34.65 | 42.46 |
| R-meas (%) | 15.2 (180.6) | 9.7 (202.6) | 7.3 (150.6) | 6.3 (173.3) | 19.8 (211.9) | 7.8 (160.1) |
| CC1/2 (%) | 99.9 (62.7) | 99.9 (56.4) | 100 (57.2) | 100 (54.6) | 99.7 (62.6) | 99.9 (70.5) |
| **Refinement statistics** | | | | | | |
| Reflections used in refinement | 18439 (1804) | 33730 (3322) | 26932 (2445) | 53091 (5230) | 24143 (2383) | 24253 (2390) |
| Reflections used for R-free | 923 (91) | 1687 (166) | 1341 (123) | 2660 (262) | 1208 (118) | 1212 (118) |
| R-work (%) | 19.30 (30.68) | 20.14 (31.17) | 23.40 (36.45) | 19.82 (37.59) | 22.29 (34.76) | 23.23 (40.97) |
| R-free (%) | 23.38 (33.29) | 23.84 (28.11) | 26.70 (40.31) | 21.76 (37.41) | 26.35 (37.31) | 26.59 (46.77) |
| Number of non-hydrogen atoms | 2417 | 2475 | 2153 | 2288 | 2370 | 2069 |
| macromolecules | 2319 | 2339 | 2047 | 2118 | 2286 | 1972 |
| ligands | 31 | 74 | 39 | 32 | 23 | 35 |
| solvent | 67 | 62 | 67 | 138 | 61 | 62 |
| Protein residues | 305 | 307 | 270 | 276 | 302 | 258 |
| r.m.s.d of bond lengths (Å) | 0.013 | 0.014 | 0.013 | 0.015 | 0.013 | 0.013 |
| r.m.s.d of bond angles (°) | 1.73 | 1.79 | 1.61 | 1.86 | 1.67 | 1.62 |
| Ramachandran favored (%) | 97.27 | 97.66 | 97.66 | 98.87 | 96.92 | 97.56 |
| Ramachandran allowed (%) | 2.73 | 2.34 | 2.34 | 1.13 | 3.08 | 2.44 |
| Ramachandran outliers (%) | 0.00 | 0.00 | 0.00 | 0.00 | 0.00 | 0.00 |
| Rotamer outliers (%) | 0.45 | 0.90 | 0.51 | 0.49 | 1.36 | 0.52 |
| Clashscore | 2.90 | 1.31 | 0.75 | 2.19 | 1.36 | 0.26 |
| Average B-factor | 48.22 | 39.00 | 43.38 | 27.93 | 45.27 | 48.45 |
| macromolecules | 48.25 | 38.61 | 42.88 | 27.28 | 45.27 | 48.38 |
| ligands | 47.86 | 48.12 | 65.64 | 36.79 | 38.39 | 47.50 |
| solvent | 47.32 | 42.83 | 45.56 | 35.85 | 47.71 | 51.00 |

**Table S2: Data collection and refinement statistics** for the crystallographic datasets of MsCI4 with bound bicyclic compounds and steroids. Values in parentheses refer to the highest resolution shell.

| | **BS700** | **BS711** | **BS713** | **Cortisone** | **Hydrocortisone hemisuccinate** |
|---|---|---|---|---|---|
| **PDB accession code** | **34BY** | **34BZ** | **34CA** | **34CB** | **34CC** |
| Wavelength (Å) | 1.000 | 1.000 | 1.000 | 1.000 | 1.000 |
| Resolution range (Å) | 37.12 -1.81 (1.88 -1.81) | 48.99 -1.9 (1.97 -1.9) | 37.26 -1.97 (2.04 -1.97) | 30.37 -2.00 (2.07 -2.00) | 48.2 -2.59 (2.68 -2.59) |
| Space group | $P2_12_12_1$ | $P2_12_12_1$ | $P2_12_12_1$ | $P2_12_12_1$ | $P2_12_12_1$ |
| a, b, c (Å) | 57.22, 58.67, 87.78 | 56.94, 59.11, 87.55 | 56.95 59.52 87.81 | 57.56, 61.14, 88.17 | 57.62, 60.8, 87.96 |
| α, β, γ (°) | 90.0, 90.0, 90.0 | 90.0, 90.0, 90.0 | 90.0, 90.0, 90.0 | 90.0, 90.0, 90.0 | 90.0, 90.0, 90.0 |
| Total reflections | 350974 (27036) | 310925 (31064) | 278862 (23816) | 284049 (26825) | 131615 (12993) |
| Unique reflections | 27505 (2701) | 23898 (2329) | 21605 (2106) | 21589 (2129) | 10081 (979) |
| Multiplicity | 12.8 (10.0) | 13.0 (13.3) | 12.9 (11.3) | 13.2 (12.6) | 13.1 (13.3) |
| Completeness (%) | 99.93 (99.67) | 99.70 (99.06) | 99.88 (99.25) | 99.90 (99.67) | 99.91 (100.00) |
| Mean I/σ (I) | 20.49 (1.14) | 11.72 (1.25) | 13.51 (1.06) | 16.51 (1.01) | 15.19 (1.56) |
| Wilson B-factor | 40.31 | 43.10 | 46.08 | 45.12 | 65.07 |
| R-meas (%) | 6.5 (179.2) | 10.2 (170.4) | 10.1 (193.1) | 10.23 (251.0) | 15.37 (181.7) |
| CC1/2 (%) | 100 (59.2) | 99.9 (81.8) | 99.9 (54.0) | 99.9 (49.7) | 99.8 (60.4) |
| **Refinement statistics** | | | | | |
| Reflections used in refinement | 27502 (2701) | 23846 (2321) | 21600 (2105) | 21583 (2128) | 10077 (979) |
| Reflections used for R-free | 1370 (133) | 1196 (113) | 1084 (109) | 1081 (106) | 503 (52) |
| R-work (%) | 21.46 (37.43) | 23.15 (39.83) | 19.97 (32.77) | 21.50 (37.52) | 25.17 (33.45) |
| R-free (%) | 24.76 (39.30) | 27.22 (39.59) | 25.37 (29.55) | 26.37 (37.10) | 30.75 (36.68) |
| Number of non-hydrogen atoms | 1777 | 2257 | 2198 | 2222 | 1832 |
| macromolecules | 1668 | 2173 | 2120 | 2109 | 1761 |
| ligands | 33 | 47 | 31 | 55 | 69 |
| solvent | 76 | 37 | 47 | 58 | 2 |
| Protein residues | 218 | 290 | 279 | 274 | 236 |
| r.m.s.d of bond lengths (Å) | 0.016 | 0.014 | 0.016 | 0.014 | 0.014 |
| r.m.s.d of bond angles (°) | 1.86 | 1.63 | 1.95 | 1.73 | 1.72 |
| Ramachandran favored (%) | 99.52 | 97.84 | 97.38 | 98.47 | 97.79 |
| Ramachandran allowed (%) | 0.48 | 2.16 | 2.62 | 1.53 | 2.21 |
| Ramachandran outliers (%) | 0.00 | 0.00 | 0.00 | 0.00 | 0.00 |
| Rotamer outliers (%) | 1.86 | 0.98 | 0.99 | 0.99 | 2.38 |
| Clashscore | 1.23 | 1.43 | 2.68 | 2.88 | 2.04 |
| Average B-factor | 45.56 | 57.29 | 57.61 | 55.44 | 63.36 |
| macromolecules | 45.19 | 57.41 | 57.59 | 55.38 | 62.50 |
| ligands | 49.05 | 53.32 | 56.51 | 56.65 | 85.81 |
| solvent | 52.05 | 55.71 | 58.97 | 56.63 | 46.83 |
